## Supplemental Figure 1 and legend for "A mitochondrial carrier transports glycolytic intermediates to link cytosolic and mitochondrial glycolysis in the human gut parasite *Blastocystis*"

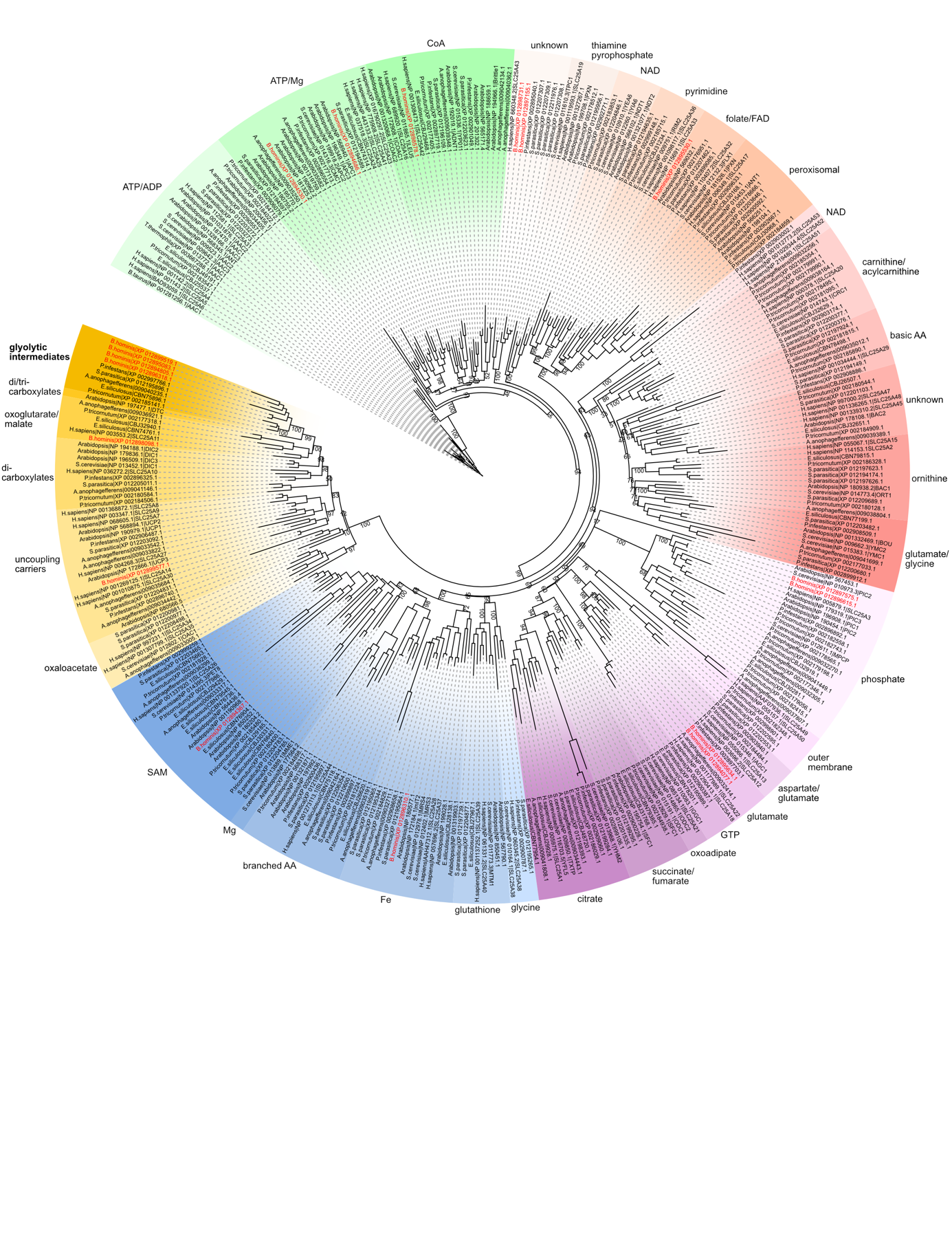

**Figure 1 - figure supplement 1**: The maximum likelihood tree was generated by IQ-TREE with the LG + F + R9 model suggested by ModelFinder. The support was calculated using ultrafast bootstrap analysis. The tree shows the relatedness of gathered human, *Saccharomyces cerevisiae*, and *Arabidopsis thaliana* mitochondrial carriers alongside a variety of stramenopile species. *Blastocystis* sequences are in red. The putative stramenopile glycolytic intermediate carriers are in dark mustard.
